# Sex chromosome and sex locus characterization in the goldfish, *Carassius auratus*

**DOI:** 10.1101/2019.12.20.875377

**Authors:** Ming Wen, Romain Feron, Qiaowei Pan, Justine Guguin, Elodie Jouanno, Amaury Herpin, Christophe Klopp, Cedric Cabau, Margot Zahm, Hugues Parrinello, Laurent Journot, Shawn M. Burgess, Yoshihiro Omori, John H. Postlethwait, Manfred Schartl, Yann Guiguen

**Author notes:** Correspondance: Yann Guiguen.

## Abstract

**Background:** Goldfish is an important model for various areas of research, including neural development and behavior and a species of significant importance in aquaculture, especially as an ornamental species. It has a male heterogametic (XX/XY) sex determination system that relies on both genetic and environmental factors, with high temperatures being able to produce female-to-male sex reversal. Little, however, is currently known on the molecular basis of genetic sex determination in this important cyprinid model. We used sequencing approaches to better characterize sex determination and sex-chromosomes in goldfish.

**Results:** Our results confirmed that sex determination in goldfish is a mix of environmental and genetic factors and that its sex determination system is male heterogametic (XX/XY). Using reduced representation (RAD-seq) and whole genome (pool-seq) approaches, we characterized sex-linked polymorphisms and developed male specific genetic markers. These male specific markers were used to distinguish sex-reversed XX neomales from XY males and to demonstrate that XX female-to-male sex reversal could even occur at a relatively low rearing temperature (18°C), for which sex reversal has been previously shown to be close to zero. We also characterized a relatively large non-recombining region (∼11.7 Mb) on goldfish linkage group 22 (LG22) that contained a high-density of male-biased genetic polymorphisms. This large LG22 region harbors 373 genes, including a single candidate as a potential master sex gene, i.e., the anti-Mullerian hormone gene (*amh*). However, no sex-linked polymorphisms were detected in the goldfish *amh* gene or its 5 kb proximal promoter sequence.

**Conclusions:** These results show that goldfish have a relatively large sex locus on LG22, which is likely the goldfish Y chromosome. The presence of a few XX males even at low temperature also suggests that other environmental factors in addition to temperature could trigger female-to-male sex reversal. Finally, we also developed sex-linked genetic markers in goldfish, which will be important for future research on sex determination and aquaculture applications in this species.

## BACKGROUND

Goldfish, *Carassius auratus*, is a domesticated fish species originating from central Asia and China that has been introduced throughout the world. Goldfish belongs to the Cyprinidae family and is considered as an important fish model for research in endocrinology [1, 2], developmental biology [3, 4] or fish pathology [5]. Thanks to the recent availability of a whole genome sequence assembly [6], goldfish is also now becoming a key model species for studies on genomics and cyprinid genome evolution. It is also a species of high aquaculture importance especially as an ornamental species, with many beautiful and sometimes bizarre phenotypes [7]. Unlike birds and mammals, sex determination in teleost is highly dynamic, with frequent turnovers of both sex determination (SD) systems [8] and master sex determining genes (MSD) [9, 10]. Currently about half a dozen different master sex determining genes have been identified in teleosts, including *dmrt1* in the Japanese medaka, *Oryzias latipes* [11], *sdY* in rainbow trout [12], *amh* in Northern pike, Nile tilapia and pejerrey [13-15], *amhr2* in yellow perch and the Takifugu pufferfish [16, 17], *gsdf* in sablefish and Luzon medaka, *O. luzonensis* [18, 19], *gsdf6a* in the turquoise killifish [20] and *sox3* in the Indian ricefish *O. dancena* [21]. MSD turnover can be evolutionarily rapid as has been shown for instance in various ricefish species [22]. In addition to genetic determinants, environmental factors -- especially temperature -- have also been shown to play a pivotal role in teleost sex determination [23]. Since the late 1960s, the goldfish sex determination system has been characterized as male heterogametic (XX/XY) [24]. More recently, a strong temperature influence on sex-ratios has also been characterized in goldfish, with high rearing temperature treatments inducing complete masculinization of chromosomally all-female genotypes (XX neomales) when applied during early 3 months development [25]. The molecular mechanisms of genetic sex determination, however, are still unknown not only in goldfish, but also in any member of the Cyprinidae family.

Because of new high throughput sequencing technologies and the availability of a whole genome sequence assembly for goldfish [26], we implemented both reduced representation (i.e., Restriction-site associated DNA sequencing (RAD-seq) [27, 28], and whole genome (i.e., Pool sequencing (Pool-seq) [29, 30]) approaches to identify sex-linked genetic polymorphisms in goldfish. We verified that identified sex-linked markers strictly segregated with the Y chromosome, and we characterized the extent of Y chromosome differentiation. Although our experiments did not identify a strong candidate sex-determining gene, these results lay a solid foundation for further molecular exploration of goldfish sex determination.

## RESULTS

### Characterization of goldfish sex-linked Y chromosome markers

Because goldfish sex determination is highly sensitive to temperature [21], with high temperature leading to the masculinization of some XX females producing XX neomales, we first searched for sex-linked markers using a RAD-seq approach that kept track of phenotypes and genotypes, potentially enabling the discrimination of XX neomales from XY genetic males. From our RAD-seq data, we identified 32 polymorphic/specific RAD-tags that were present in 12-15 males among the 30 phenotypic males used in this experiment, and completely absent in all the 30 phenotypic females (Fig. 1A, Supplementary excel file1). These results suggest a male heterogametic genetic sex determination system (XX/XY) as previously shown in goldfish [24], but with a rather high occurrence of XX neomales (around 50 %) in this population of two-year old animals raised outdoor and obtained from different batches of animals with different spawning times i.e., from May-June to late September.

**Figure 1.**
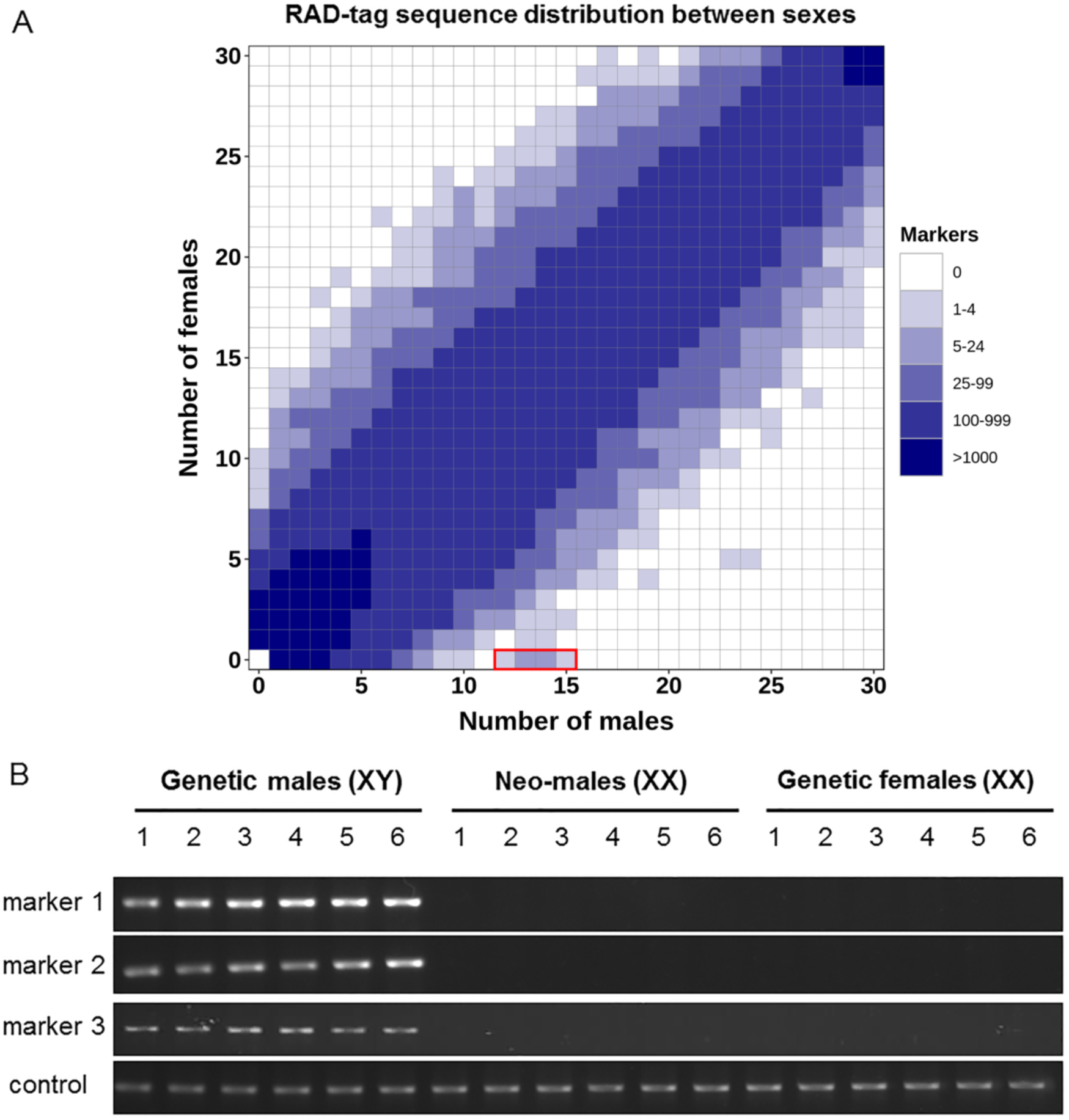
RAD-sex tags and male-specific markers in goldfish. (**A**) Haplotypes heatmap in phenotypic males and females’ goldfish. Each cell in the heatmap represents the number of haplotypes presented in x phenotypic males and y phenotypic females (x: cumulative number of males, y: cumulative number of females). Haplotypes present in more than 12 males and absent in all females were identified as male-specific haplotypes (highlighted by red box). (**B**) Genotyping of goldfish males and females with three Y-allele primer pairs and one autosomal primer pair used as a positive control. Goldfish are categorized into three groups i.e., putative genetic males (XY), putative XX neomales, and genetic females by combining the results of both Y-allele genotyping and sex phenotyping.

To validate the hypothesis that these markers were linked to the heterogametic sex (XY) and the Y chromosome, we first sequenced using Illumina reads and assembled a draft genome sequence of a male goldfish identified as a putative XY male based on the polymorphic/specific RAD-tags (see Material & Methods) and blasted these 32 marker sequences against this genome assembly. This analysis returned 20 contigs with highly significant matches (Supplementary excel file2) spanning a total of 0.24 Mb. By anchoring these sex-linked RAD sequences on our genome assembly, we were able to design three putative Y-allele specific primer pairs that were used to genotype the same individual animals that were used for the RAD-seq analysis. PCR genotyping using these three primer pairs accurately discriminated putative XY genetic males from putative XX neomales and females (Fig. 1B), validating that these primers accurately identified the two types of males found in our RAD-seq analysis. We then genotyped male breeders from our experimental stock with these primers and selected one putative XX neomale (breeder 1, negative PCR amplifications) and one putative XY male (breeder 2, positive PCR amplifications); and both individuals were crossed to the same XX female to generate two separate batches of fish. If our Y-allele specific primers correctly identify the Y chromosome, then our putative XX neomale should give only female offspring and the putative XY male should give both male and female offspring. These two experimental populations were then reared at low temperature (18°C) during the first three months after fertilization to minimize high temperature masculinization [25], and were subsequently maintained at 24°C for nine additional months before the identification of the phenotypic sex. Results from the histological examination of the offspring gonads of the putative XX neomale identified 7 fish with testes, 83 fish with ovaries, and 41 fish with undifferentiated gonads. Disregarding fish with undifferentiated gonads suggests a sex ratio of 7.8% males and 92.2% females for the offspring of the XX neomale (Table 1). Gonadal histology of the offspring of the putative XY revealed 48 animals with testes, 65 with ovaries, and 14 with undifferentiated gonads, which gives a sex ratio of 42.5% male and 57.5% female, ignoring the offspring with undifferentiated gonads (Table 1). These sex ratio differences (Table 1), strongly support the hypothesis that male breeder 1 is an XX neomale with an offspring sex ratio not significantly different from an expected all-female population with a slight percentage of female-to-male sex-reversal, and that breeder 2 is a genetic XY male with an offspring sex ratio not significantly different from an expected 50:50 sex ratio. In agreement with these results, none of the XX neomale offspring produced a positive PCR amplification for our three Y-allele specific primer pairs (Figure S3, Table 2), and all 48 phenotypic males but only one of 65 phenotypic females offspring from the XY phenotypic male produced positive amplifications (Figure S4, Table 2). This result also indicates that no neomales were detected in offspring from the XY genetic male if we do not consider the undetermined individuals compared to the 7.8% of neomales in the XX neomale offspring population.

**Table 1.**
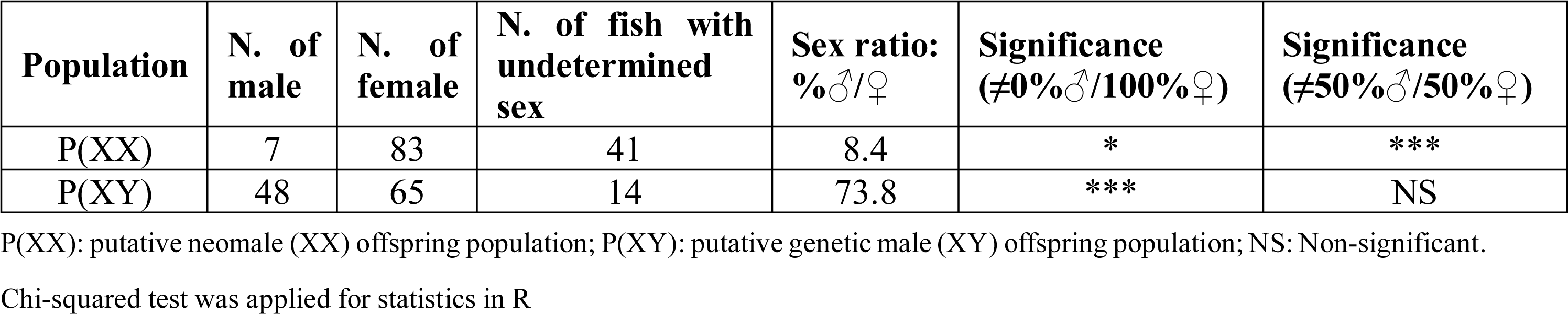
Statistics of phenotypic sex in two populations.

**Table 2.** Goldfish Y-allele sex-linkage.

| Population | Male<sup>#</sup> | Female<sup>#</sup> | Undetermined sex<sup>#</sup> | Sex linkage |
| --- | --- | --- | --- | --- |
| P(XX) | 0 / 7 | 0 / 83 | 0 / 41 | NS |
| P(XY) | 48 / 48 | 1 / 65 | 10 / 14 | *** |

<sup>#</sup> Y-allele positive genotyping / total number of samples. P(XX): putative neomale (XX) offspring population; P(XY): putative genetic male (XY) offspring population; NS: Non-significant. Fisher's exact test was applied for statistics in R

### Characterization of the goldfish sex chromosome and sex-determining region (SDR)

Using the three Y-allele specific primer pairs described above, we genotyped goldfish individuals and selected 30 phenotypic and genotypic males that were used along with 30 phenotypic females to contrast whole genome sex differences by pool-sequencing analysis [29]. Pool-sequencing reads from the respective XY male and phenotypic female pools were mapped to the high contiguity goldfish female genome assembly [6] to characterize genomic regions enriched for sex-biased signals, i.e., sex coverage differences or sex-biased Single Nucleotide Polymorphism (SNP) differences. Whole genome analysis of SNP distribution (Figure 2) revealed a strong sex-linked signal in males on linkage group 22 (LG22) and two unplaced scaffolds (NW_020523543.1 and NW_020523609.1) with a high density of observed SNPs being heterozygous in the male pool and homozygous in the female pool (Y-specific allele). Interestingly, of the 32 markers found using the RAD-Seq approach, 7 tags were enriched in the unplaced scaffold NW_020523543.1 (Fig. 3C), confirming by a second approach that this scaffold is part of the SD locus in goldfish. These regions with a high density of male-specific SNPs (Figure 3) are potential sex-determining regions that could contain the goldfish master sex determining gene. LG22, being the only linkage group with a large sex determining region (SDR, highlighted by a black box on Fig. 3A, C, D) containing a high-density of male-specific SNPs (∼11.7 Mb), likely corresponds to the goldfish Y sex chromosome.

**Figure 2.**
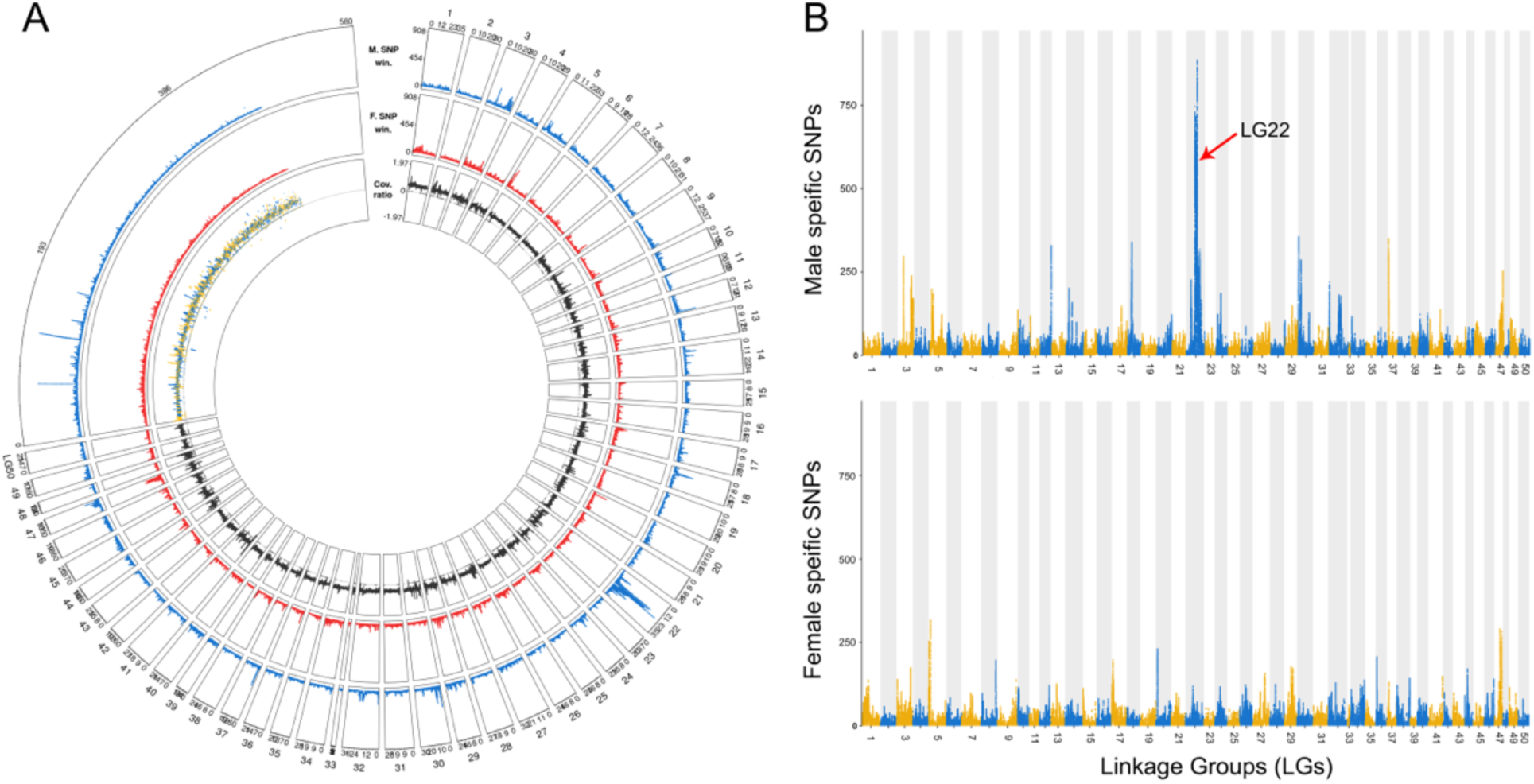
Sex determining regions identified by remapping the Pool-seq male and female reads onto the female genome assembly. SNPs were counted using 100kb sliding window with an output point every 500bp. (**A**) Circular plot showing the genome wide metrics of the Pool-seq analysis. All the 50 goldfish linkage groups (LGs) are labelled with their LG number and all unplaced scaffolds are fused together. Outer to inner tracks show respectively: the male-specific SNPs, the female-specific SNPs, and the reads depth ratio between males and females. (**B**) Manhattan plot of the male- and female-specific SNPs showing a strong enrichment of male-specific SNPs on LG22.

**Figure 3.**
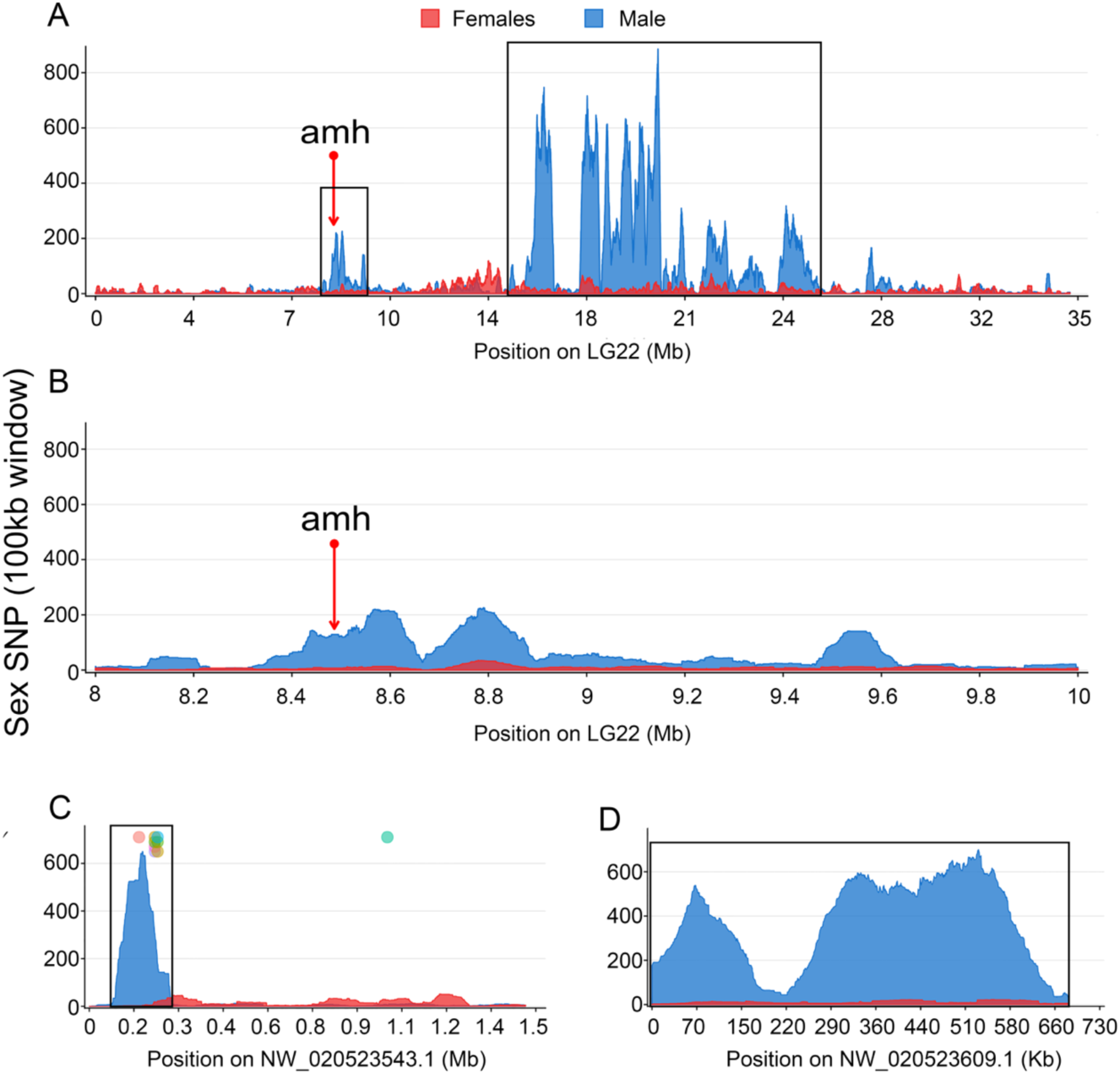
Distribution of male-specific SNPs on LG22 and unplaced scaffolds NW_020523543.1 and NW_020523609.1. SNPs were counted using 100kb sliding window with an output point every 500bp and female- and male-specific SNPs were respectively indicated by red and blue color. (**A**) A large sex-determination region was identified on LG22, which is highlighted with a black box. The candidate sex-determining gene *amh* is located on this LG22, but not in the high density, male-specific SNP region. The region from 8Mb to 10Mb containing *amh* is zoomed in panel (**B**). (**C**) The NW_020523543.1 unplaced scaffold exhibits a region around 0.1Mb harboring a small region (200 kb) with a high-density of male-specific SNPs. Meanwhile, sequence comparisons demonstrate that 7 male-biased RAD-tags (colored circles) on a total of 32 map with a high identity onto this scaffold. In contrast, few female-specific SNPs were enriched on this scaffold (red area). (**D**) The unplaced NW_020523609.1 scaffold is enriched in male-specific SNPs.

We also observed, however, some smaller signals with less dense sex-linked SNPs in other linkage groups (Figure 2A) like for instance on LG47 (Fig. S1) with both male and female sex-linked signals. Interestingly, LG47 is paralogous to LG22 stemming from the Cyprinidae whole genome duplication [6]. Indeed, due to this recent common ancestry, these two chromosomes share large homologous and syntenic regions (Fig. S2) that could have resulted in some false remapping of the pool-sequencing reads leading to some of these secondary minor signals.

### Identification of candidate SD genes

Searches for annotated genes by BLAST within the 20 contigs found in our male goldfish draft genome assembly based on the RAD-Seq approach did not return any matches for a candidate SD gene, but mostly transposable elements (Supplementary excel file 3). All genes within the SDR (N= 373) were extracted because they are potential candidates for being SD gene(s) (Supplemental excel file 4). Interestingly, among these genes the anti-Mullerian hormone gene (*amh*) was found at the beginning of the SDR on LG22 (Fig. 3B). This gene has been reported to be a sex-determining gene in other fish species [14, 15]. However, we did not identify any male-specific SNPs in the coding sequence of goldfish *amh*. In addition, other male specific alleles within the 5kb promoter region did not show any sex-linkage.

## DISCUSSION

Though goldfish is an important economic ornamental fish and a useful model for studying development, evolution, neuroscience, and human disease [3], characterization of goldfish sex-specific sequences and potential sex chromosomes have not been reported. In this study, we explored goldfish sex determination using two complementary whole-genome approaches and found that this species has a XX/XY sex determination system as previously described [24] and a large, non-recombining sex determination region on LG22. Although RAD-sequencing or pool-sequencing have been often used separately to explore sex determination in vertebrates [16, 30, 31], we choose to combine these two approaches in goldfish because of the significant female-to-male sex reversal induced by temperature [25] that would have prevented a clear identification of the sex determining region using only a pooled strategy, which mixes genetic XY males and XX males resulting from the sex reversal of genetic females. Because RAD-sequencing keeps track of each individual, we were able to identify sex-reversed individuals in goldfish that might have masked sex-linked markers in Pool-seq.

Sex markers identification is an important step to characterize SD systems [32-38]. Using two complementary whole-genome approaches, we characterized genomic regions containing sex-linked markers. In goldfish, these sex-linked markers are genomic DNA variations including gaps, indels and SNPs that present heterozygote polymorphisms in all males and complete homozygosity in all females. This male-specific heterozygosity pattern agrees with a male heterogametic XX/XY system as previously reported using progeny testing of hormonally sex-reversed breeders [24]. We found, however, a strong environmental influence leading to a relatively high proportion (around 50%) of female-to-male sex-reversal in the first experimental population that we used for the RAD-Sequencing approach. These animals were actually two-year old goldfish raised in an outdoor experimental facility and obtained at different spawning times i.e., from May-June to late September. Some of these animals experienced early development during summer time at potentially higher temperature and others had their early developmental period at lower temperatures. Considering the known effects of high temperature on female-to-male sex reversal in goldfish [25], the fact that some of these fish were exposed to a high summer temperature could explain this relatively high percentage of female-to-male sex-reversed animals. This high percentage was not found in our other experiments in which fish were raised in indoor recirculating system facilities with a tightly controlled low temperature (18°C) maintained throughout the whole early development phase (3 months). This situation indeed confirms earlier findings showing that temperature is probably a major trigger of neomasculinization in goldfish, but we also found that even at this low temperature there was still a small percentage of female-to-male sex-reversal (7.8%), suggesting that other environmental factors, potentially social factors as demonstrated in other species [8, 39], could also play a role on goldfish sex determination. Apart from goldfish, sex determination in other teleost fish, including Tilapia [40], medaka [41] and tongue sole [42] is also regulated by temperature, which overrides the genetic sex determination mechanisms and leads to female-to-male sex reversal. By developing genetic sexing tools in goldfish that allows the identification of Y-allele carrying animals, we also brought additional evidence that some of these phenotypic males were indeed sex-reversed XX genetic females. These genetic sexing tools are indeed important for better deciphering genetic and environmental sex determination in goldfish. But these PCR primers could be also used to facilitate the industrial production of commercial goldfish-related hybrid fish in China [43, 44], by helping to identify neomales i.e., XX female-to-male sex reversed animals.

Sex determination in vertebrates is highly variable with the major exceptions of Eutherian mammals and birds in which XX/XY and ZZ/ZW monofactorial sex determination systems have been conserved over a long evolutionary period [45, 46]. In contrast, fish exhibit much more diverse and dynamic sex determination [9, 10, 47], with monofactorial and polyfactorial [48, 49] genetic systems and frequent switches and turnovers of master sex-determining genes [12, 14, 15, 17, 21, 50]. In goldfish, we identified male-specific markers and obvious male-specific SNPs strongly enriched on LG22. This result confirms that goldfish has an XX-XY system [24] and also indicated that LG22 is the sex chromosome in that species. Evidence is accumulating for the hypothesis that sex chromosomes, in most cases, evolve from autosomes with *de novo* initial evolution of a new sex determination mechanism that subsequently becomes fixed and extended by the suppression of recombination on the sex chromosome in the vicinity of the initial sex locus, which may increase the size of this non recombining sex determination locus [51]. In goldfish, ∼11.7 Mb of LG22 contains numerous male-specific SNPs. A similar large size of the non-recombining region on the sex chromosomes was also found in tilapia including 17.9 Mb in *Sarotherodon melanotheron* and 10.7 Mb in *Oreochromis niloticus* [30, 31]. The large non-recombining region on LG22 contains 373 gene models based on the goldfish genome annotation and also a large number of transposable elements (TEs) that were found to be strongly enriched in the male specific contigs identified by our RAD-Sex and our draft genome analysis. Enrichment of TEs around sex loci has been found in other vertebrate species [52] and may play a crucial role for suppression of recombination leading to an expansion of sex chromosome divergence.

With LG22 being the potential sex chromosome in goldfish, it is reasonable to believe that the non-recombining region that we characterize on LG22 contains the goldfish master sex determining gene. But the only “usual suspect” master sex determining candidate found in this region and the additional non-assembled scaffolds containing sex-linked markers is the anti-Mullerian hormone gene (*amh*) that is located at the beginning of the LG22 non-recombining region. Duplications of *amh* have been characterized as the master sex determining gene in different fish species [14, 15], making Amh and members of the TGF-beta pathway [17, 19, 20] likely candidates for this sex-determining function. But we have not been able to characterize sex-linked variation neither in the *amh* coding DNA sequence nor in its 5 kb proximal promoter sequence. Even if we cannot rule out the hypothesis that *amh* regulation could be affected by sex-specific cis-regulatory elements located very far upstream from *amh*, our results do not provide any clear and direct evidence that this gene is the goldfish master sex determining gene. Indeed, not all master sex determining genes are classical “usual suspects” known to be involved in the sex-differentiation pathway like TGF-beta members [17, 19, 53], Sox3 [21], or Dmrt1[50, 54]. For instance, the rainbow trout master sex determining gene arose from the duplication / transposition / evolution of an immune-related gene [12]. This finding suggests that goldfish could also have an unusual master sex determining gene, preventing an easy and direct identification just with simple genome-wide analyses and candidate gene approaches.

The goldfish genome, like the genomes of the common carp and other species of the cyprinid subfamily cyprininae is characterized by a relatively recent whole genome duplication (WGD) that occurred approximately 14 million years ago [6]. This WGD adds an extra complexity to our search for sex-linked regions and sex determining candidate genes because some of these duplicated regions may still retain large blocks of high sequence similarity. The cyprininae genome duplication probably explains why we found an additional sex-biased signal on LG47 that stems from the duplication of the same ancestral chromosome that LG22. In addition to the cyprininae WGD, the current goldfish reference genome sequence [6] was assembled from the sequences of an XX gynogenetic animal, meaning that the LG22 sex chromosome sequence is an X chromosome sequence in which potential Y specific regions may be not present. We indeed produced a first draft genome sequence of an XY male but a higher contiguity male genome including long-read technology would be needed to better explore sex-chromosome differences and characterize potential sex-determining candidates.

## CONCLUSIONS

Our results confirm that sex determination in goldfish is a complex mix of environmental and genetic factors, and that its genetic sex determination system is male heterogametic (XX/XY). We also characterized a relatively large non-recombining region (∼11.7 Mb) on LG22 that is likely to be the goldfish Y chromosome. This large non-recombining region on LG22 contains a single obvious candidate as a potential master sex gene, namely the anti-Mullerian hormone gene (*amh*). No sex-linked polymorphism, however, was detected in the goldfish *amh* gene and its 5 kb proximal promoter sequence. Our work provides the foundation required for additional studies that are now required to better characterize sex determination in goldfish and to characterize its master sex-determining gene.

## MATERIALS AND METHODS

### Experiment fish

Fish used for RAD-seq and Pool-seq were reared outdoors and obtained from different spawning times i.e., between May-June and late September. Putative XY and XX males were selected using Y-allele specific primers and these two males were crossed with the same female to produce two goldfish populations that were incubated and reared indoor at 18°C during three months after fertilization to minimize the chance of sex reversal induced by temperature according to previous research [25]. After these 3 months at 18°C, the rearing temperature was gradually increased to 24°C over a period of 7 days to avoid suddenly dramatic temperature variation. One-year old fish were euthanized with Tricaine before dissection. Gonads of goldfish were fixed in Bouin’s fixative solution for 24 hours and then embedded gonads were cut serially into 7 µm sections and stained with Hematoxylin to characterize ovarian or testicular features. Fin clips were stored in 90% alcohol for DNA extraction and genotyping. Statistics were applied to test for significant sex ratio differences and genotype/phenotype sex-linkage with a Chi-squared test (p < 0.05).

### DNA extraction and genotyping

For genotyping, fin clips were lysed with 5% Chelex and 20 mg Proteinase K at 55°C for 2 hours, and subsequently denatured by Proteinase K at 99°C for 2 min. Supernatant containing genomic DNA (gDNA) was collected to a new tube after a brief centrifugation. Finally, DNA was diluted to half and stored at −20°C. For genome sequencing, gDNA was extracted with NucleoSpin Kits for Tissue (Macherey-Nagel, Duren, Germany) following the manufacturer’s instructions. gDNA concentration and quality were measured with a NanoDrop ND2000 spectrophotometer (Thermo Scientific, Wilmington, DE) and a Qubit3 fluorometer (Invitrogen, Carlsbad, CA).

Primers were designed from the sequences of male-biased contigs for sex genotyping and a positive control (Table S1) based on contig flattened_line_0 from our Illumina male genome assembly (Accession number: WSJC00000000) using Primer3 version 0.4.0 (http://primer3.ut.ee). PCRs were performed with 0.1 µM of each primer, 50 ng of gDNA adjusted at 50 ng/µl, 100 µM dNTP mixture, and 1 µl of 10× PCR Buffer (Sigma Aldrich) with 0.25 units of JumpStart Taq DNA Polymerase (Sigma Aldrich) in a total volume of 25 µl. The PCR thermal cycle procedures were: 94°C for 30s for denaturing, 58°C for 30s for annealing and 72°C for 30s for extending for 35 cycles. Finally, PCR products were electrophoresed on 1.5% agarose gels.

### Restriction-site association sequencing (RAD-seq) and male-marker discovery

Genomic DNA was extracted from 30 males and 30 females and digest with the restriction enzyme *SbfI* for constructing a RAD-seq library according to standard protocols [55]. Briefly, for each sample, 1µg of DNA was digested using *SbfI*. Digested DNA was purified using AMPure PX magnetic beads (Beckman Coulters) and ligated to indexed P1 adapters (one index per sample) using concentrated T4 DNA ligase (NEB). Ligated DNA was purified using AMPure XP magnetic beads. Each sample was quantified using microfluorimetry (Qubit dsDNA HS assay kit, Thermofisher) and all samples were pooled in equal amount. The pool was fragmented on a Biorputor (Diagenode) and purified using a Minelute column (Qiagen). Sonicated DNA was size selected on an 1,5 % agarose cassette aiming for an insert size of 300 bp to 500 bp. Size selected DNA was extracted from the gel using the Qiaquick gel extraction kit (Qiagen), repaired using the End-It DNA-end repair kit (Tebu Bio) and adenylated on its 3’ ends using Klenow (exo-) (Tebu-Bio). P2 adapter was ligated using concentrated T4 DNA ligase (NEB) and 50 ng of the ligated product was engaged in a 12 cycles PCR. After AMPure XP beads purification, the resulting library was checked on a Bioanalyzer (Agilent) using the DNA 1000 kit and quantified by qPCR using the KAPA Library quantification kit (Roche, ref. KK4824). The library was sequenced on one lane of Hiseq2500 in single read 100nt mode using the clustering and SBS v3 kit following the manufacturer’s instructions.

Raw reads were demultiplexed with the program *process_radtags.pl* of Stacks with default settings. 135,019,110 (79.1%) reads were kept after this procedure. Demultiplexed reads were subsequently processed by the RADSex software version 2.0.0 (http://github.com/RomainFeron/RadSex). The distribution of sequences between male and female were calculated with function *distrib* with all settings to default. This distribution of sequences was visualized with *plot_sex_distribution* function of radsex-vis (http://github.com/RomainFeron/RADSex-vis) (Fig 1.A). Sequences significantly associated with sex were extracted using the function *signif*, which identifies sex-bias tags. Male-biased tags were compared to the male *de novo* assembly with ncbi-blast+ (version: 2.6.0) setting the e-value cutoff to 1^e-20^ to identify long, homologous male-biased contigs. Male specific PCR primers were designed from these contigs sequences (see Table S1) using Primer3 version 0.4.0 (http://primer3.ut.ee).

### Pooled genome sequencing (Pool-seq) and sex differentiated region identification

Genomic DNA extracted from the fin clips of 13 phenotypic females and 13 genotypic males selected from the animals used for the RAD-Seq experiment, were used for the Pool-Seq analysis. The 13 genotypic males were genotyped using the three Y-allele PCR primers described above. Genomic DNA were pooled in equimolar ratio according to sex and Pool-seq libraries were generated using the Truseq nano DNA sample prep kit (Illumina, ref. FC-121-4001) following the manufacturer’s instructions. Briefly, each pool was sonicated using a Bioruptor (Diagenode). The sonicated pools were repaired, size selected on magnetic beads aiming for a 550 pb insert size and adenylated on their 3’ ends. Adenylated DNA was ligated to Illumina’s specific adapters and, after purification on magnetic beads, was amplified in an 8 cycles PCR. Libraries were purified using magnetic beads, checked on a Fragment Analyzer (Agilent) using the HS NGS Fragment kit (DNF-474-33) and quantified by qPCR using the KAPA Library quantification kit (Roche, ref. KK4824). Each library was sequenced on half a lane of a rapid v2 flow cell (Illumina) in paired end 2×250nt mode.

Reads from the male and female pools were remapped to a genome sequence coming from a gynogenesis-derived female [QPKE00000000] using BWA mem version 0.7.17 with default parameters. Then, BAM files were sorted and merged with Picard tools version 2.18.2 with default parameters. After that, PCR duplicates were removed with Picard tools. Reads with mapping quality less than 20 and that did not map uniquely were also removed with Samtools version 1.8. Subsequently, the two sex BAM files were used to generate a pileup file using samtools mpileup with per-base alignment quality disabled (-B). A sync file was created using popoolation mpileup2sync version 1.201 (parameters: --min-qual 20), which contains the nucleotide composition of each sex for each position in the reference. Finally, with this sync file, SNPs and coverage between the two sexes of all reference positions were overall calculated with PSASS (version 2.0.0, doi:10.5281/zenodo.2615936). We used a 100kb sliding window with an output point every 500bp to identify sex-specific SNPs enriched regions with PSASS. The PSASS parameters were as follows: minimum depth set to 10 (--min-depth 10), range of heterozygous SNP frequency for a sex-linked locus 0.5±0.2 (--freq-het 0.5, --range-het 0.2), homologous SNP frequency for a sex-linked locus >0.98 (--freq-hom 1, --range-hom 0.02), overlapped sliding window (--window-size 100000, --output-resolution 500). Data visualization was implemented with an R package (http://github.com/RomainFeron/PSASS-vis).

### Sequencing and *de novo* assembly of a goldfish male genome

One genetic male was selected for *de novo* assembly using the Y-specific primers described above. Library was generated using the Truseq nano DNA sample prep kit (Illumina, ref. FC-121-4001) following the manufacturer’s instructions. Briefly, DNA from a single male individual was sonicated using a Bioruptor (Diagenode). The sonicated DNA was repaired, size selected on magnetic beads aiming for a 550 pb insert size and adenylated on its 3’ ends. Adenylated DNA was ligated to Illumina’s specific adapters and, after purification on magnetic beads, was amplified in an 8 cycles PCR. Library was purified using magnetic beads, checked on a Fragment Analyzer (Agilent) using the HS NGS Fragment kit (DNF-474-33) and quantified by qPCR using the KAPA Library quantification kit (Roche, ref. KK4824). The library was sequenced on one lane of a rapid v2 flow cell (Illumina) in paired end 2*250nt mode. Illumina paired-end reads were assembled using DiscovarDeNovo (reference https://software.broadinstitute.org/software/discovar/blog/) with standard parameters.

## ABBREVIATIONS

RAD-seq: Restriction site-associated DNA sequencing;
SNP: Single nucleotide polymorphism;
SD: Sex determination;
SDR: Sex differentiated region,
MSD: master sex determining genes.

## DECLARATIONS

### Ethics approval

Research involving animal experimentation conformed to the principles for the use and care of laboratory animals, in compliance with French (“National Council for Animal Experimentation” of the French Ministry of Higher Education and Research and the Ministry of Food, Agriculture, and Forest) and European (European Communities Council Directive 2010/63/UE) guidelines on animal welfare.

### Consent for publication

Not applicable

### Availability of data and material

This Whole Genome Shotgun project has been deposited at DDBJ/ENA/GenBank under the accession WSJC00000000. The version described in this paper is version WSJC01000000. Genome sequencing reads of the male genome, the male and female pool-sequencing reads and the RAD-seq demultiplexed sequences have been deposited in the Sequence Read Archive (SRA), under BioProject PRJNA592334.

### Competing interests

The authors declare that they have no competing interests.

### Funding

This project was supported by funds from the “Agence Nationale de la Recherche” and the “Deutsche Forschungsgemeinschaft” (ANR/DFG, PhyloSex project, 2014-2016) to MS and YG. Montpellier Genomics (MGX) facility was supported by France Génomique National infrastructure, funded as part of “Investissement d’avenir” program managed by Agence Nationale pour la Recherche (contract ANR-10-INBS-09). This work was also supported by Grant-in-Aid for Scientific Research (19K22426) to YO, and grants R01OD011116 and 5R01GM085318 from the USA National Institutes of Health to JHP.

## Authors’ contributions

Conceived and designed the experiments: YG, MW

Funding acquisition: YG, MS, JP, LJ

Investigation: MW, MP, JG, EJ, AH, CR, HP, SB, YO

Bioinformatics analysis: RF, CK, CC, MZ

Visualization: MW

Wrote the paper: MW, YG

### Acknowledgements

We are grateful to the Genotoul bioinformatics platform Toulouse Midi-Pyrenees (Bioinfo Genotoul) for providing computing and/or storage resources and to the INRA-LPGP experimental facilities for taking care of goldfish experiments.

## Supplementary information

**Supplementary excel file 1:** Sequences of putative Y-allele RAD-tags (N= 32) found in some males but absent from all females.

**Supplementary excel file 2:** Contigs from a goldfish Illumina male genome assembly with homologies with the putative Y-allele RAD-tags.

**Supplementary excel file 3:** Annotation of potential Y chromosome contigs by sequence comparisons to NCBI Non-redundant protein sequence database using blastx.

**Supplementary excel file 4:** Detailed information of annotated genes in the goldfish sex determination regions extracted from the NCBI genome annotation file (accession number QPKE00000000).

---

**Table S1.**
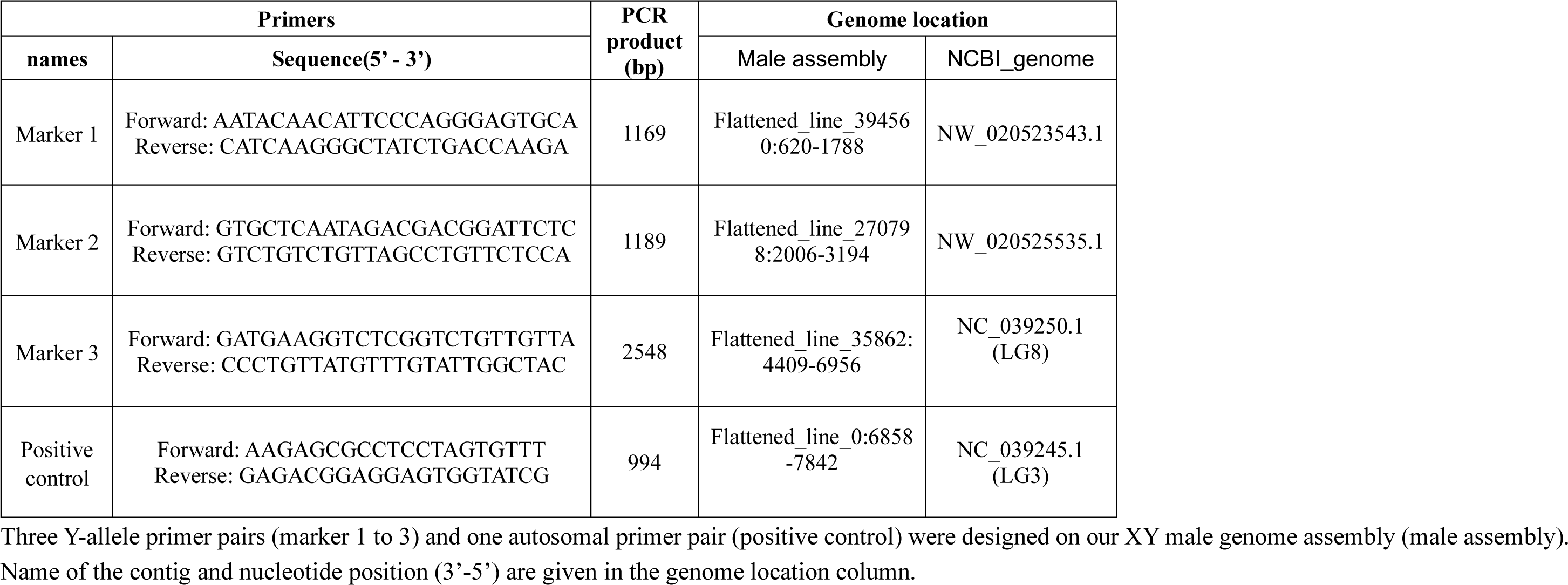
Sequences of the primers used for Y-allele genotyping in goldfish.

## SUPPLEMENTARY FIGURES

**Figure S1:**
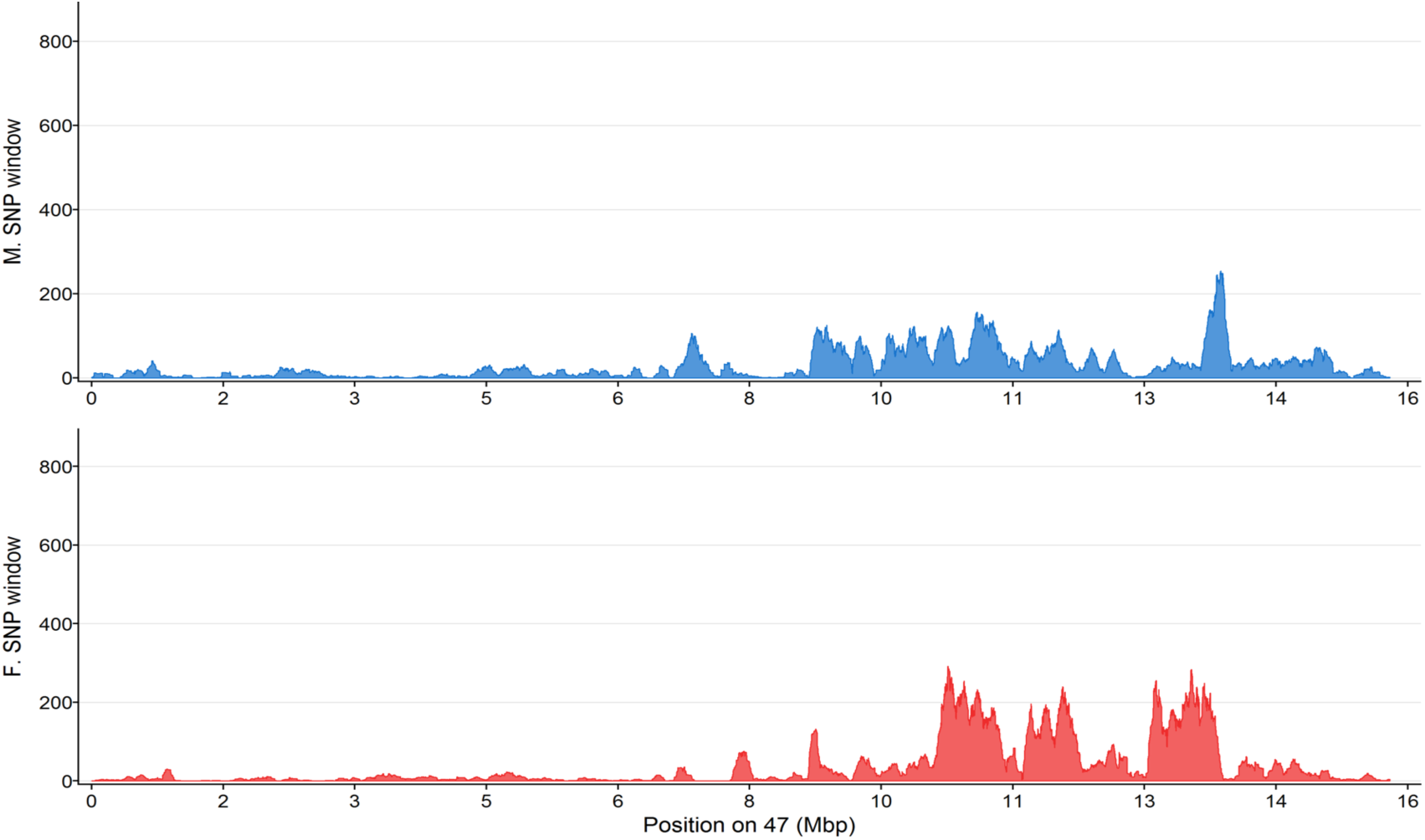
Distribution of sex-biased SNPs on LG47. SNPs were counted using 100kb sliding window with an output point every 500bp. The top panel displays the profile of male-specific SNPs (blue area), while the bottom panel displays the profile of female-specific SNPs (red area).

**Figure S2:**
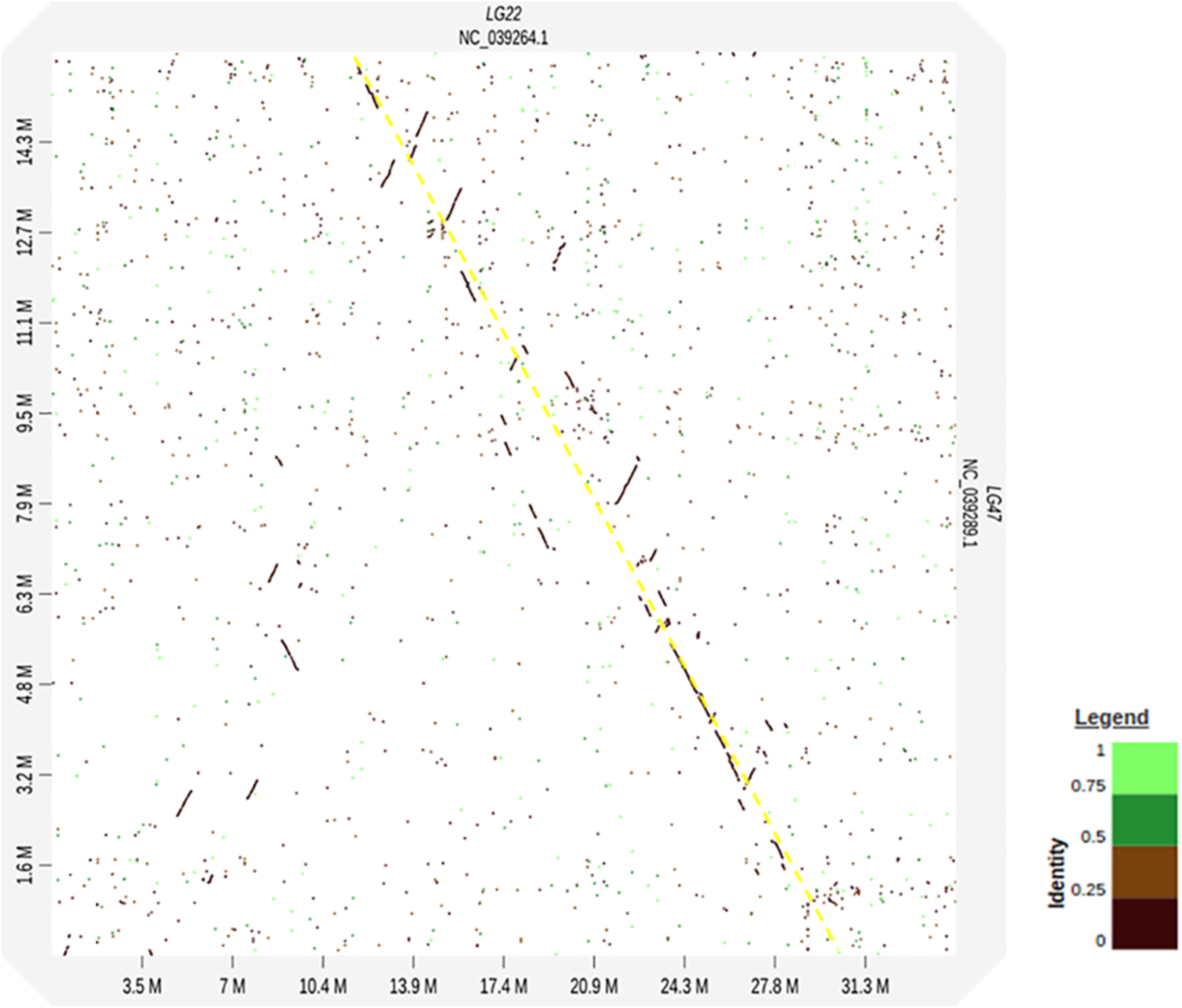
Dot plot comparison of LG22 and LG47 showing conserved synteny between these two linkage groups.

**Figure S3:**
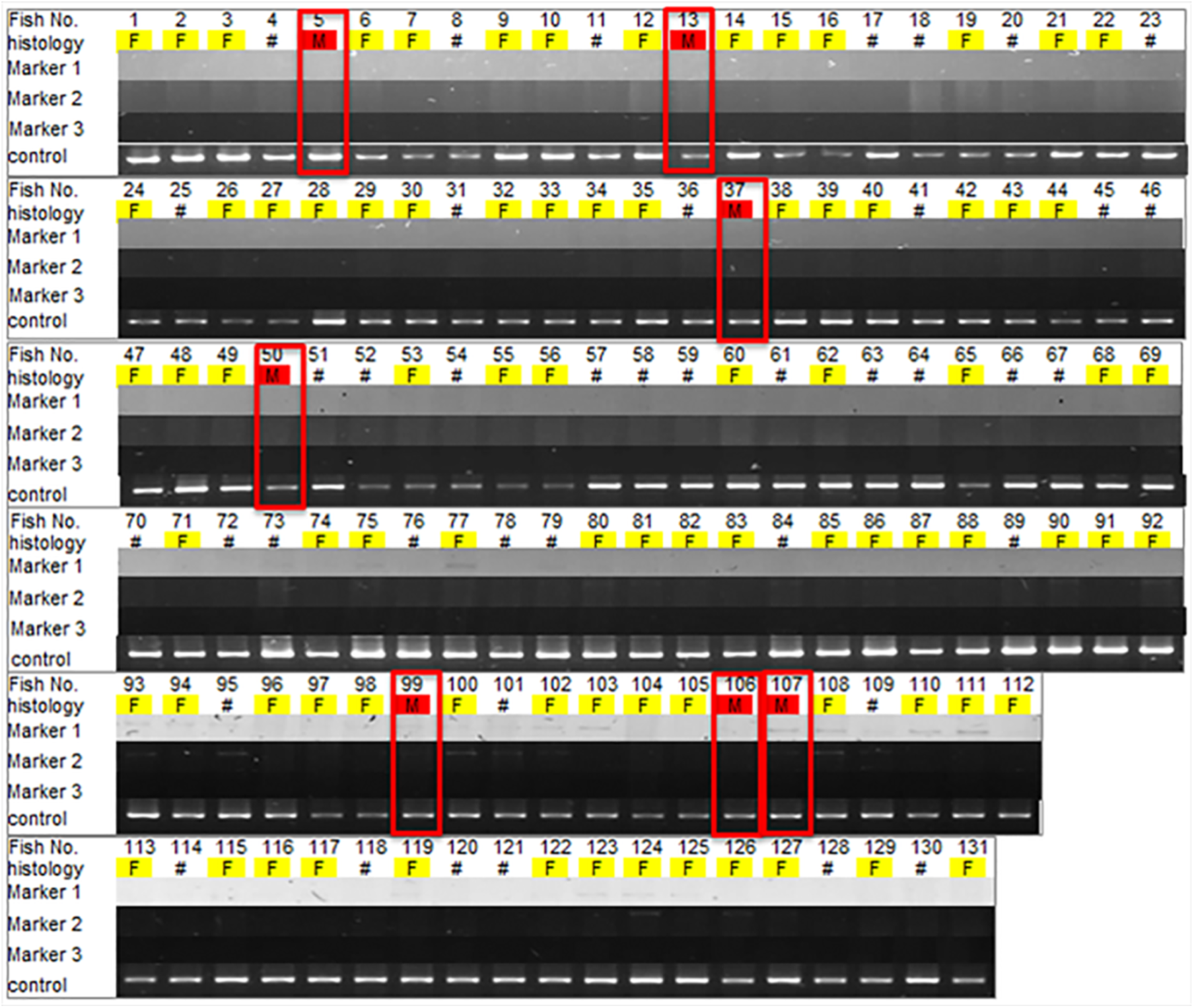
Sex genotyping with Y-allele primers of the offspring of a putative XX neomale with a normal XX female. Genotyping was conducted with three Y-allele primers and one autosomal primer used as a gDNA quality control. Phenotypic sex was determined by gonadal histology and males and females are shown using red and yellow color respectively. Female-to-male sex-reversed animals (N= 7) are highlighted by red boxes. Hashes indicate animals with unknown phenotypic sex with undifferentiated gonads based on histology.

**Figure S4.**
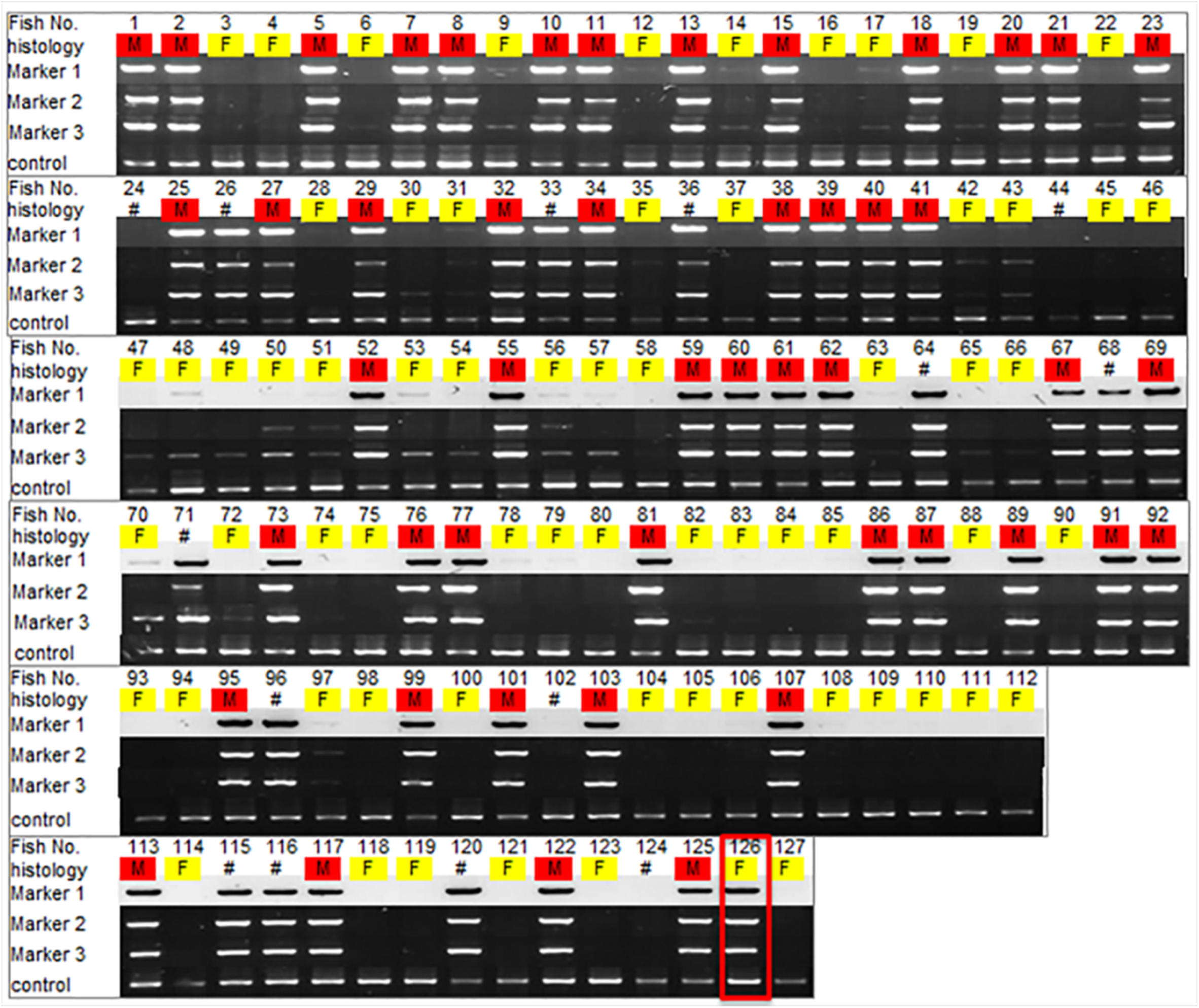
Sex genotyping with Y-allele primers of the offspring of a putative XY male with a normal XX female. Genotyping was conducted with three Y-allele primers and one autosomal primer used as a gDNA quality control. Phenotypic sex was determined by gonadal histology and males and females are shown using red and yellow color respectively. The female-to-male sex-reversed animal (N= 1) is highlighted by a red box. Hashes indicate animals with unknown phenotypic sex with undifferentiated gonads based on histology.

